# Ticks; a reservoir for virus emergence at the human-livestock interface in Uganda

**DOI:** 10.1101/2023.03.10.532017

**Authors:** Stella A. Atim, Shirin Ashraf, Marc Niebel, Alfred Ssekagiri, Maryam N. Hardy, James G. Shepherd, Lily Tong, Anna R Ademun, Patrick Vudriko, Joseph Erume, Steven Odongo, Denis Muhanguzi, Willy Nguma, Teddy Nakayiki Dip, Joyce Namulondo, Ana Filipe, Julius J Lutwama, Pontiano Kaleebu, Charles Masembe, Robert Tweyongyere, Emma C. Thomson

## Abstract

**Background:** Uganda is one of the most biodiverse regions on the planet and a hotspot for virus emergence. In particular, the warm-humid lowlands favour tick population growth with the associated risk of tick-borne disease. The prevalent tick species *Rhipicephalus appendiculatus, R. evertsi evertsi* and *Amblyomma variegatum* harbour a diverse range of viruses, from harmless to highly pathogenic. Notably, the orthonairoviruses cause human outbreaks of Crimean-Congo haemorrhagic fever (CCHF) regularly within the cattle corridor of Uganda, a region spanning from the south-west to the north-east of the country.

**Methods:** In the ArboViral Infection (AVI) study, the first to explore the virome of ticks in Uganda using next generation sequencing (NGS), we collected ticks from three geographically diverse areas and subjected these to target-enrichment (TE) NGS. Viral genomes were detected by *de novo* assembly, mapping and BLASTn.

**Results:** We analyzed a total of 2,754 ticks collected from 31 livestock farms in the districts of Arua, Nakaseke and Lyantonde. These were combined into 219 pools by site of collection and tick species, including *R. appendiculatus, R. evertsi evertsi*, *A. variegatum* and *Hyalomma rufipes*. We detected partial or near-complete viral genomes in 163 tick pools; 110 (67%) of which were from Arua, 39 (24%) from Nakaseke and 12 (7%) from Lyantonde districts. 2 pools (2%) were from Arua/Lyantonde. These included 22 species of virus, representing 15 genera and 9 families, including the *Nairoviridae*, *Retroviridae*, *Orthomyxoviridae*, *Chuviridae*, *Rhabdoviridae*, *Phenuiviridae, Parvoviridae, Poxviridae* and *Flaviviridae*. There were 8 viral species known to be pathogens of humans or animals and 5 highly divergent genomes detected, representing novel virus species. A high abundance of orthonairoviruses was notable, including CCHFV, Dugbe virus and a novel *Orthonairovirus* species that we have named Macira virus.

**Interpretation:** Ticks in Uganda are an important reservoir of diverse virus species, many of which remain uncharacterised and of unknown pathogenic potential.

**Author Summary:** Ticks are parasitic arachnids that may transmit a spectrum of viral diseases to humans and animals. Uganda is a hotspot for such tick-borne diseases. In this study, we sequenced ticks collected from three geographically diverse regions of Uganda using a semi-agnostic next-generation sequencing method in order to detect viruses from all known virus families. We collected and analyzed 2,754 ticks from 31 farms across the country. Within these ticks, we detected 22 species of virus from 15 genera and 9 viral families, including 8 animal or human pathogens and 5 new novel virus species. Notably, orthonairoviruses, including the highly pathogenic Crimean-Congo haemorrhagic fever virus, were highly prevalent in the ticks. The researchers suggest that ticks in Uganda serve as an important reservoir for diverse viruses, many of which have significant pathogenic potential. This information will inform public health efforts to prevent and control tick-borne diseases in Uganda and other similar regions.

## Introduction

Ticks are obligate hematophagous arthropods that parasitise fish, amphibians, reptiles, birds, and mammals in every geographic region on the planet. (1-3). There are over 900 tick species classified into two main families; *Ixodidae* (hard ticks) and *Argasidae* (soft ticks), and a third family, *Nuttalliellidae*, with only one species, *Nuttalliella namaqua*, exhibiting features associated with both hard and soft ticks (4). Over ten percent of tick species vector pathogens, including viruses, bacteria, protozoa and helminths. Viral pathogens within the *Asfarviridae*, *Reoviridae*, *Rhabdoviridae*, *Orthomyxoviridae*, *Peribunyaviridae*, *Nairoviridae* and *Flaviviridae* are particularly well-represented (5, 6). These may be amplified in wild mammalian hosts and livestock with the associated risk of pathogen spill over into the human population (7). In Uganda, Crimean-Congo haemorrhagic fever virus (CCHFV) frequently causes outbreaks of devastating and often fatal disease with significant public health implications (8, 9).

The viral diversity in ticks in Uganda and the surrounding Central and East African regions has been previously only partially described. The emergence of next generation sequencing (NGS) technologies provides the opportunity to describe the eco-epidemiology of the tick virome to help direct public health policy for vector and pathogen control. In the ArboVirus Infection (AVI) study, we aimed to determine virus diversity of *Ixodidae* ticks parasitizing livestock to improve understanding of tick-borne viruses circulating in central and north- western Uganda to ascertain the associated risk of arboviral infection in surrounding animal and human populations.

## Results

We analyzed a total of 2754 ticks collected from 31 farms in the districts of Arua (n=13), Nakaseke (n=12) and Lyantonde (n=6) (**Fig 1, Supplementary Table 1**). The ticks were combined into 219 pools, including pools of *R. appendiculatus (*n=107)*, R. evertsi evertsi* (n=46), *A. variegatum* (n=35), including adults, nymphs and larval pools (**Table 1**).

**Fig 1:**
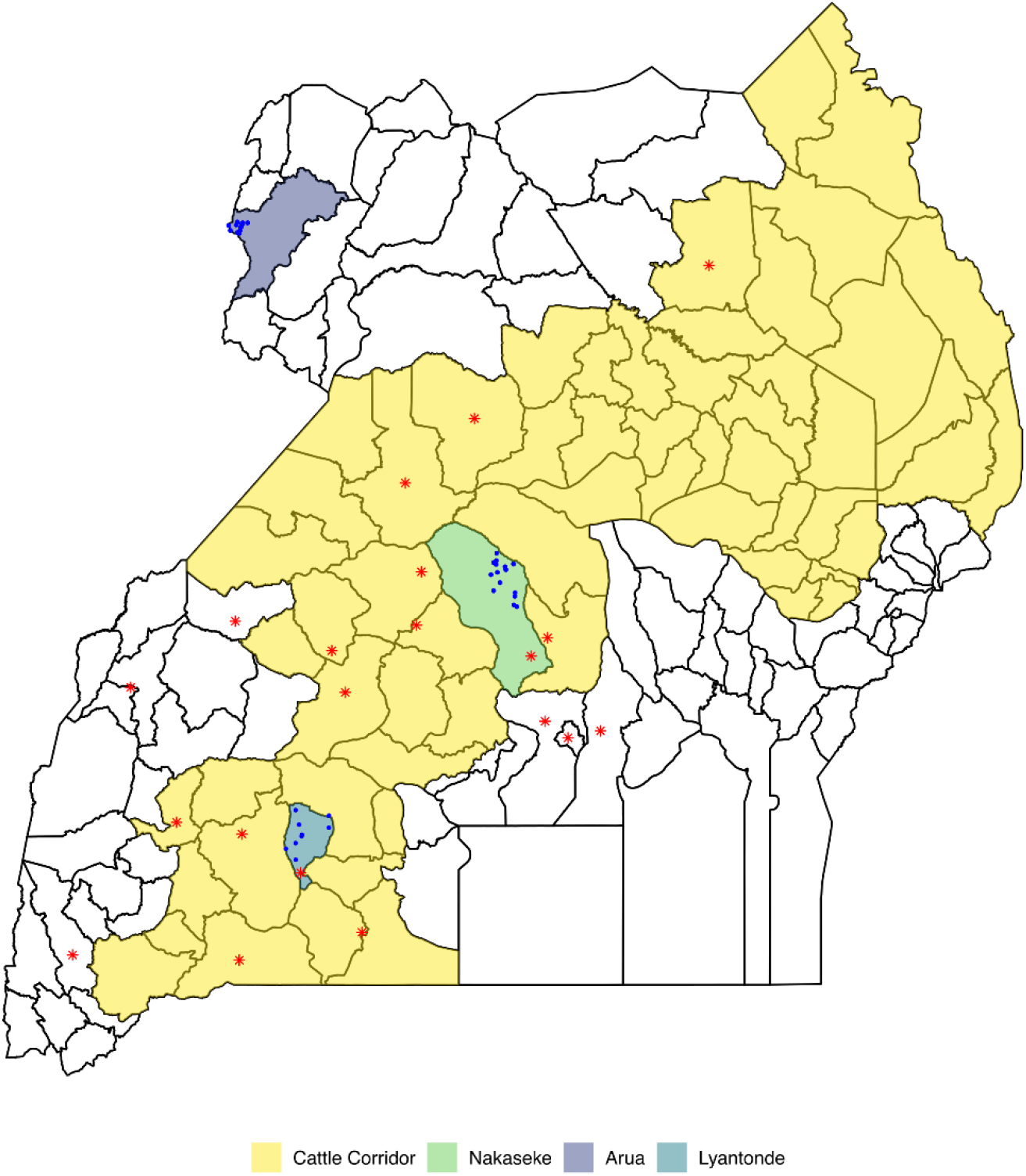
Map of Uganda showing selected study districts. The blue dots represent sampled farms, the yellow belt is Uganda’s cattle corridor inclusive of Lyantonde and Nakaseke districts, and the red asterisks are locations of CCHF human outbreaks between 2013-2019.

**Table 1:** Ticks analyzed by target enrichment NGS.

| Tick species | No. of ticks by host |  |  |  | Total no. of ticks | No. of pools |
| --- | --- | --- | --- | --- | --- | --- |
|  | Cattle | Goats | Dogs | Environment |  |  |
| *R. appendiculatus* | 1239 | 151 | 5 | 13 | 1408 | 107 |
| *R. evertsi evertsi* | 200 | 286 | - | - | 486 | 46 |
| *R. boophilus decoloratus* | 26 | - | - | - | 26 | 2 |
| *R. boophilus microplus* | 41 | - | - | - | 41 | 7 |
| *A. variegatum* | 388 | - | - | - | 388 | 35 |
| *Hyalomma rufipes* | 1 | - | - | - | 1 | 1 |
| *Rhipicephalus* nymphs | 31 | 115 | - | 115 | 261 | 13 |
| *Amblyomma* nymphs | 118 | - | - | - | 118 | 7 |
| *Amblyomma* larvae | - | - | - | 25 | 25 | 1 |
| Totals | 2044 | 552 | 5 | 153 | 2754 | 219 |

Viruses were identified in 163 tick pools, 110 of which were from Arua, 12 from Lyantonde and 39 from Nakaseke districts (**Fig 2**). 2 pools were from Arua and Lyantonde. Of the virus-positive tick pools, 139 derived from *Rhipicephalus* ticks and 29 were from *Amblyomma* ticks. Viruses detected belonged to 9 families (*Nairoviridae, Retroviridae, Orthomyxoviridae, Chuviridae, Rhabdoviridae, Phenuiviridae, Parvoviridae, Poxviridae and Flaviviridae*), 15 genera (*Alpharicinrhavirus, Bandavirus, Betaretrovirus, Ephemerovirus, Erythroparvovirus*, *Flavivirus, Jingmenvirus, Ledantevirus, Mivirus, Orthonairovirus, Parapoxvirus, Pestivirus, Phlebovirus, Quaranjavirus, Thogotovirus*) and 22 species including 5 novel virus species (**Table 2; Supplementary Table 2, 3**). 68 contigs of <1kb in length were classified by family or genus only and are likely to represent novel species that could not be fully speciated according to International Committee on Taxonomy of Viruses (ICTV) criteria.

**Fig 2:**
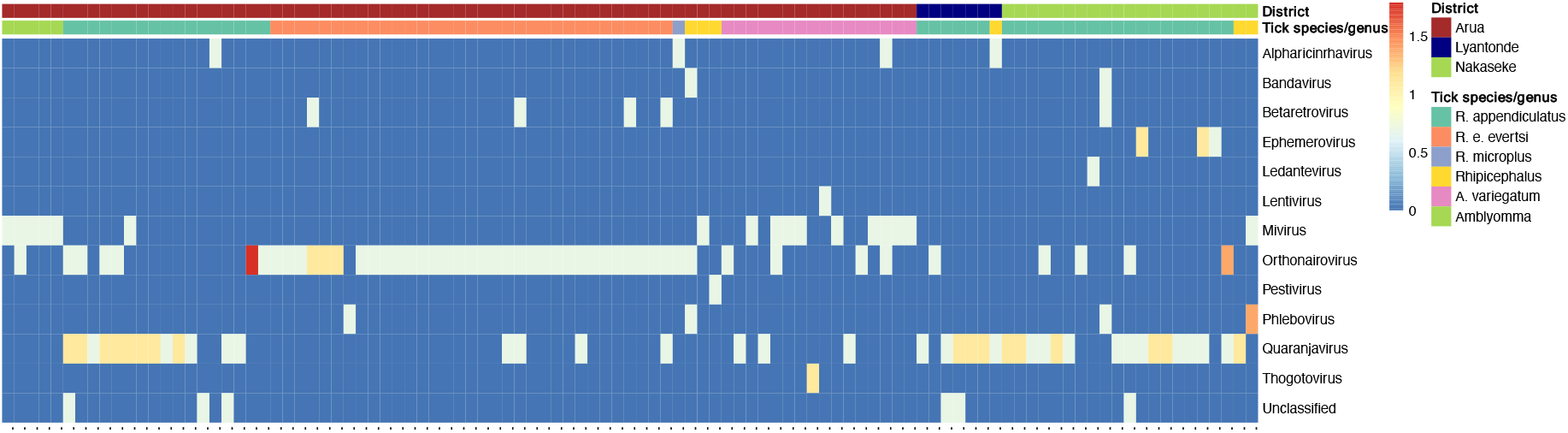
Heatmap showing the abundance of selected virus genera across sampling districts and tick species. Individual tick pools are shown on the horizontal axis and virus genera on the vertical axis. The abundance was log transformed and visualized using the pheatmap package in R.

**Table 2:** Virus species detected in ticks, by tick source and species

| Virus species | Cattle |  | Cattle<br>(total) | Dog<br>*Rhip.* | Dog<br>(total) | Env.<br>*Rhip.* | Env.<br>(total) | Goat<br>*Rhip.* | Goat<br>(total) | Species<br>(total) |
| --- | --- | --- | --- | --- | --- | --- | --- | --- | --- | --- |
|  | *Amb.* | *Rhip.* |  |  |  |  |  |  |  |  |
| ***Chuviridae*** | **14** | **3** | **17** |  |  |  |  | **1** | **1** | **18** |
| Karukera tick virus | 13 | 2 | 15 |  |  |  |  | 1 | 1 | 16 |
| Unclassified | 1 | 1 | 2 |  |  |  |  |  |  | 2 |
| ***Flaviviridae*** |  | **12** | **12** |  |  | **3** | **3** | **6** | **6** | **21** |
| Bole tick virus 4 |  | 1 | 1 |  |  | 2 | 2 |  |  | 3 |
| Engoha virus (new) |  | 2 | 2 |  |  |  |  | 2 | 2 | 4 |
| Jingmen tick virus |  | 3 | 3 |  |  |  |  | 1 | 1 | 4 |
| Trinbago virus |  | 2 | 2 |  |  | 1 | 1 | 3 | 3 | 6 |
| Unclassified |  | 4 | 4 |  |  |  |  |  |  | 4 |
| ***Nairoviridae*** | **6** | **26** | **32** |  |  | **2** | **2** | **19** | **19** | **53** |
| CCHF virus | 1 | 4 | 5 |  |  |  |  | 1 | 1 | 6 |
| Dugbe virus | 5 | 7 | 12 |  |  |  |  | 4 | 4 | 16 |
| Macira virus (new) |  |  |  |  |  | 1 | 1 |  |  | 1 |
| Nairobi sheep<br>disease virus |  |  |  |  |  |  |  | 1 | 1 | 1 |
| Unclassified |  | 15 | 15 |  |  | 1 | 1 | 13 | 13 | 29 |
| ***Orthomyxoviridae*** | **4** | **31** | **35** |  |  |  |  | **9** | **9** | **44** |
| Thogoto<br>thogotovirus | 1 |  | 1 |  |  |  |  |  |  | 1 |
| Unclassified | 3 | 15 | 18 |  |  |  |  | 6 | 6 | 24 |
| Zambezi tick virus 1 |  | 16 | 16 |  |  |  |  | 3 | 3 | 19 |
| ***Parvoviridae*** | **1** |  | **1** |  |  |  |  |  |  | **1** |
| Bovine parvovirus 3 | 1 |  | 1 |  |  |  |  |  |  | 1 |
| ***Phenuiviridae*** |  | **1** | **1** |  |  |  |  | **4** | **4** | **5** |
| Enkuda virus (new) |  | 1 | 1 |  |  |  |  |  |  | 1 |
| Enkwa virus (new) |  |  |  |  |  |  |  | 1 | 1 | 1 |
| Forecariah virus |  |  |  |  |  |  |  | 1 | 1 | 1 |
| Palma virus |  |  |  |  |  |  |  | 1 | 1 | 1 |
| Torwonik virus<br>(new) |  |  |  |  |  |  |  | 1 | 1 | 1 |
| ***Poxviridae*** | **1** |  | **1** |  |  |  |  |  |  | **1** |
| Pseudocowpoxvirus | 1 |  | 1 |  |  |  |  |  |  | 1 |
| ***Retroviridae*** | **1** |  | **1** |  |  |  |  | **5** | **5** | **6** |
| Enzootic nasal<br>tumor virus of goats |  |  |  |  |  |  |  | 5 | 5 | 5 |
| Unclassified | 1 |  | 1 |  |  |  |  |  |  | 1 |
| ***Rhabdoviridae*** | **1** | **8** | **9** | **1** | **1** |  |  | **2** | **2** | **12** |
| Barur virus |  | 1 | 1 |  |  |  |  |  |  | 1 |
| Rhipicephalus<br>associated rhabdo-like virus |  | 2 | 2 |  |  |  |  |  |  | 2 |
| Unclassified |  | 4 | 4 | 1 | 1 |  |  | 1 | 1 | 6 |
| Wuhan Tick Virus 1 | 1 | 1 | 2 |  |  |  |  | 1 | 1 | 3 |
| **Unclassified** | **1** | **1** | **2** |  |  |  |  |  |  | **2** |
| Unclassified | 1 | 1 | 2 |  |  |  |  |  |  | 2 |
| Total | 29 | 82 | 111 | 1 | 1 | 5 | 5 | 46 | 46 | 163 |

The median virus genus richness ranged between 1 and 2 and Simpson’s diversity ranged between 0 and 0.5 among pools in all districts and tick species without significant overall differences between districts. The most abundant genera were *Orthonairovirus* and *Quaranjavirus.* Quaranjaviruses were abundant in *R*. *appendiculatus* tick pools across all three districts. The proportion of *Quaranjavirus* positive pools was significantly higher in both Nakaseke (p=0.001) and Lyantonde (p=0.02) versus Arua district. We also observed a high abundance of orthonairoviruses in *R*. *evertsi* tick pools from Arua district. The proportion of *Orthonairovirus* positive pools was significantly higher in Arua district (p=0.003). (**Supplementary Table 4**).

For each sample and virus species, the longest virus contig was placed onto a pre-existing amino acid alignment using an evolutionary placement algorithm (EPA-NG)(10). This method cannot classify new viruses but places contigs onto phylogenetic branches based on regions of homology in order to illustrate the diversity of non-overlapping contigs across the cladogram of each virus family (**Fig 3**). The *Nairoviridae* family cladogram (**Fig 3a**) is represented by a single genus; *Orthonairovirus*. Most contigs were placed terminally, notably within the CCHFV species clades (with human (red tip) and mammal (blue tip) host ranges). Within the *Phenuiviridae* family (**Fig 3b**), placements were made within the *Bandavirus, Phlebovirus* and *Uukuvirus* genera. Notable species placements included the human pathogen Bhanja virus and Forecariah virus (purple), an arthropod-restricted species. Four genera were identified within the *Rhaboviridae* (**Fig 3c**) with contig placements in the *Alpharicinrhavirus, Ledantevirus* and *Ephemerovirus* genera. Both the *Orthomyxoviridae* PB1 (**Fig 3d**) and *Flaviviridae* (**Fig 3e**) were represented with placements within the *Quaranjavirus* and *Pestivirus* genera respectively. Virus contigs were also detected within the segmented *Flaviviridae* (**Fig 3f**). Other viral families identified included by phylogenetic placement included within the *Retroviridae* gag protein, *Chuviridae* polymerase and *Dicistroviridae* capsid protein (**S1 Fig**).

**Fig 3:**
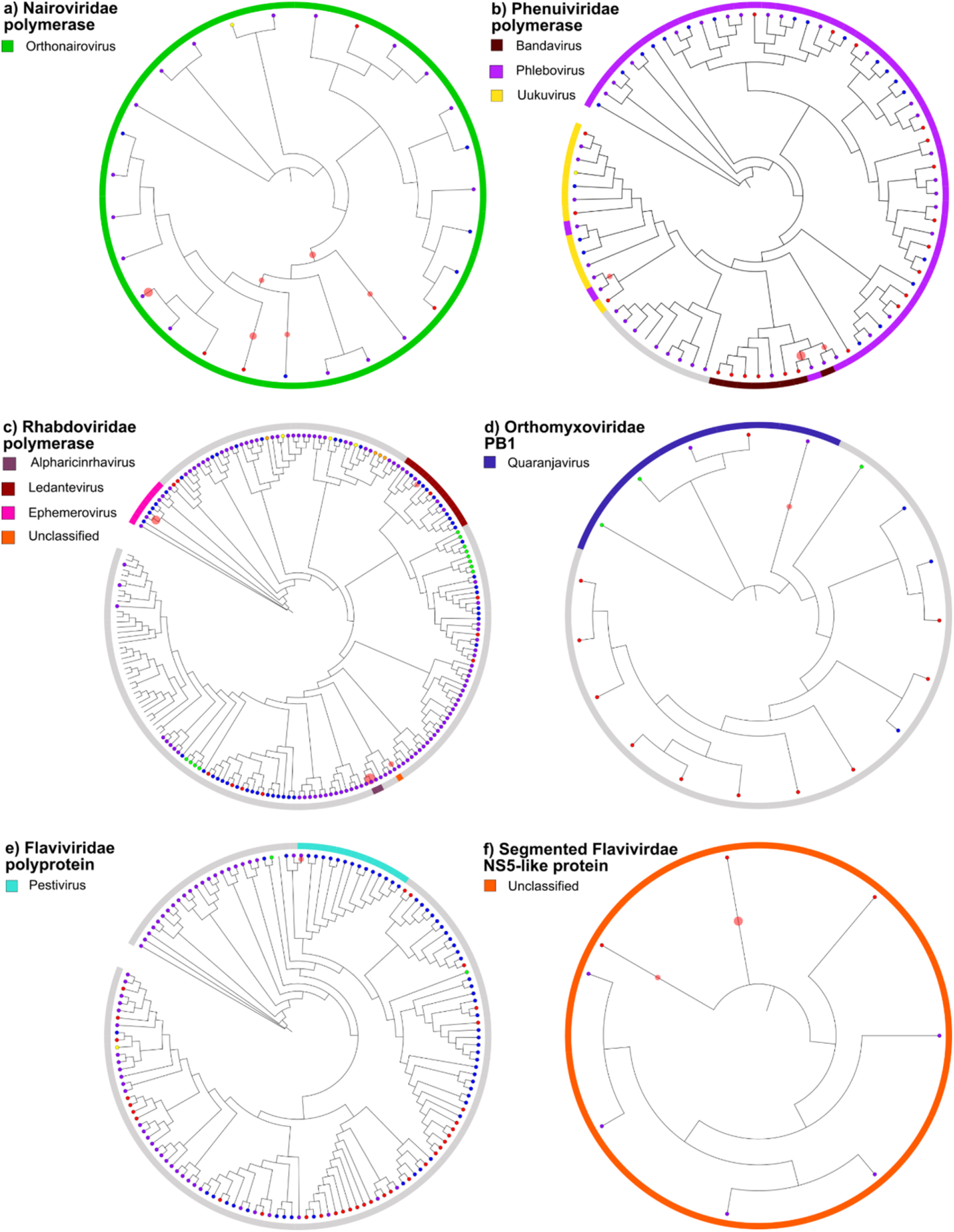
Phylogenetic placement cladogram illustrating the diversity of viruses detected in ticks across multiple virus families. EPA-NG phylogenetic trees were generated using reference trees constructed using IQTREE to illustrate the diversity of non-overlapping viral amino acid contigs within different families. The number of samples with a contig sitting on each branch is proportionate to the size of the red circles. Branch tips are labelled as red (human), blue (mammal), green (fish), yellow (bird), orange (reptile), purple (arthropod). Genera are represented by the outer circle (colours as labelled in key legends).

Next, maximum likelihood phylogenetic analysis of viral genome sequences was carried out to further characterise circulating viral species in ticks. We restricted this analysis to contigs of >1kb in order to generate reliable phylogenetic data. Viral genomes within the *Nairoviridae*, *Orthomyxoviridae*, segmented *Flaviviridae, Flaviviridae, Rhabdoviridae, Chuviviridae* and *Phenuiviridae* were subjected to maximum likelihood phylogenetic analysis with relevant reference sequences (**Fig 4; S2 Fig)**. Five of these genomes represent novel virus species: Macira virus, Engoha virus, Torwonik virus, Enkwa virus, and Ekuda virus.

**Fig 4:**
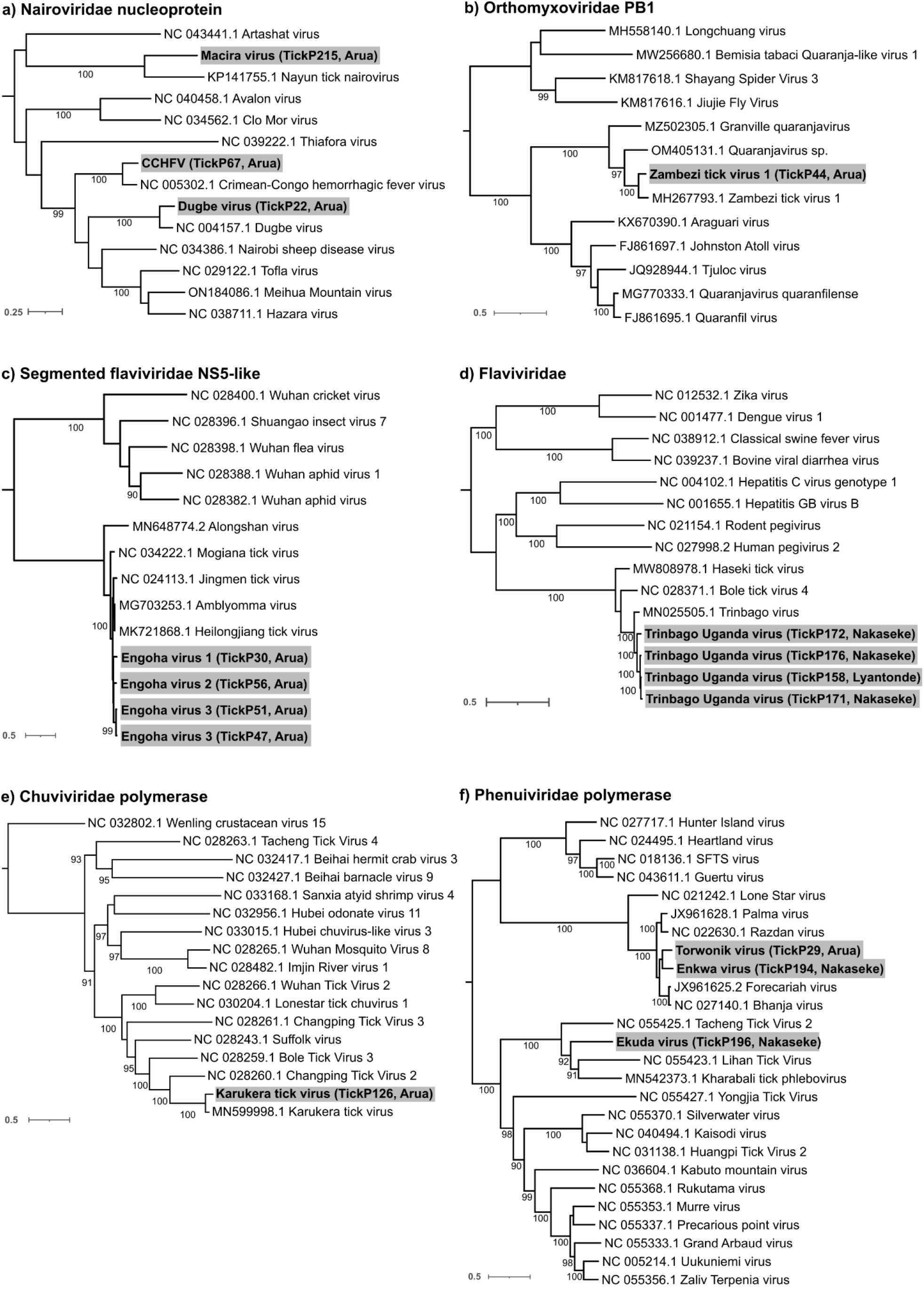
Maximum likelihood phylogenetic analysis of viral nucleotide sequences. Representative sequences derived from the **a** *Nairoviridae* N protein (P215 represents a novel species; P67 is one of four closely related sequences and P22 is one of four closely related sequences), **b** *Orthomyxoviridae* PB1 protein **c** Segmented *Flaviviridae* NS5-like protein, **d** *Flaviviridae* polyprotein **e** *Chuviviridae* polymerase protein and **f** *Phenuiviridae* polymerase protein. Samples are indicated in bold and highlighted in grey. The scale for each tree is shown in the bottom left corner of each tree. Bootstrap values based on 1000 replicates are shown on the branches.

Fifty-three pools were found to contain contigs that mapped within the family *Nairoviridae*. Five independent *R. appendiculatus* and one *Amblyomma* tick pools contained CCHFV genome. Four of these (P9, P67, P143, P171) when mapped using the closest available references had sufficient genome across each of the L, M and S segments to carry out maximum likelihood phylogenetic analysis (**Fig 5**). While the L and S segments of these clustered with other sequences in the Clade II (Africa 2) lineage, there was evidence of reassortment of the M segment in two cases (P67 and P143) in different localities, with a highly divergent M segment, most closely related to viruses from the Group V (Europe I) geographic lineage. We also detected Dugbe virus (DUGV) in 16 tick pools of *Rhipicephalus* and *Amblyomma* from Arua region, of which 8 were used for phylogenetic analysis (P8, P17, P18, P19, P21, P22, P24, P26). A representative sequence from pool P22 derived from *A. variegatum* ticks in Arua district is shown in **Fig 4a**.

**Fig 5:**
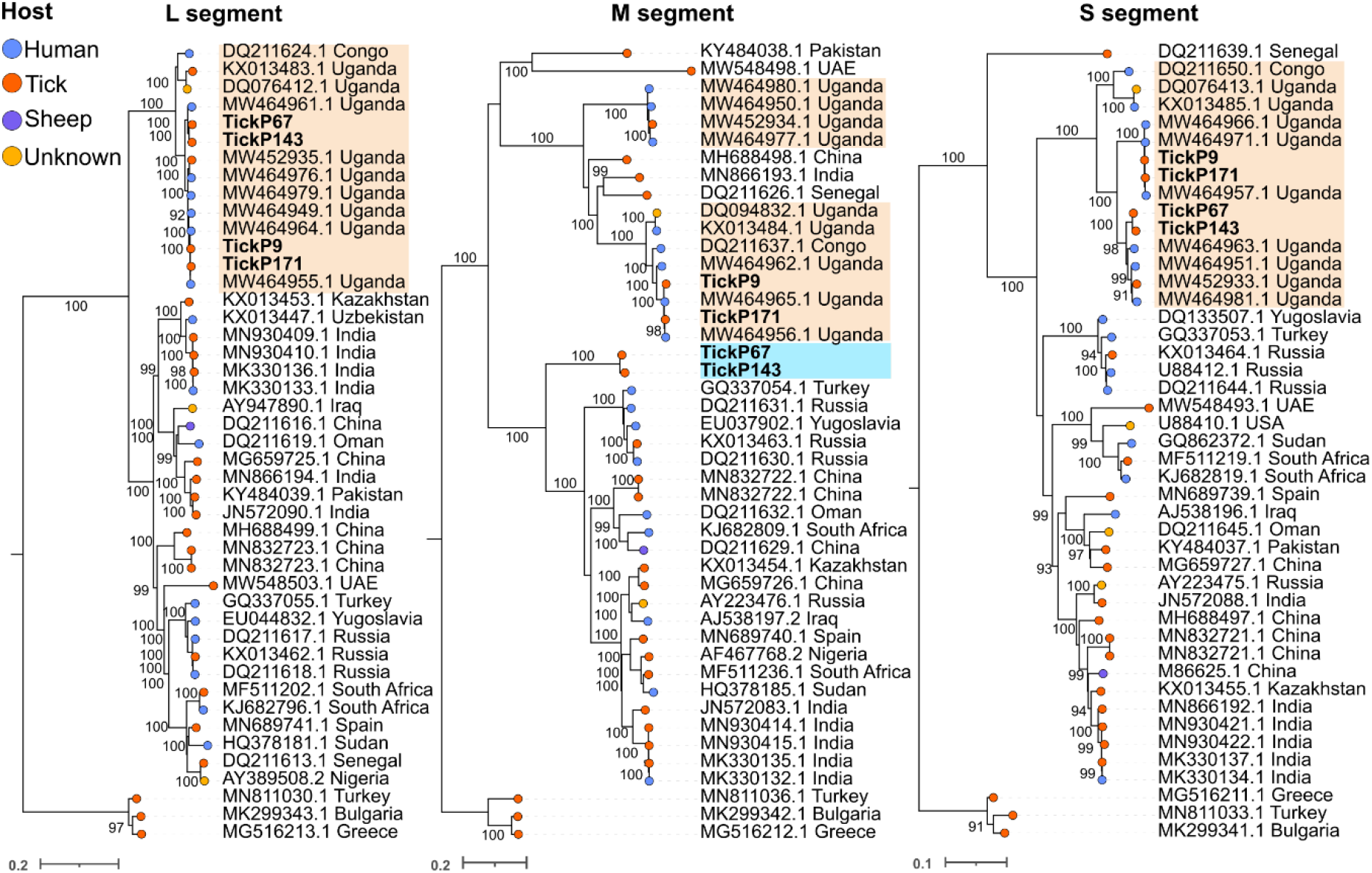
Maximum likelihood phylogenetic analysis of CCHFV sequences. Samples obtained from *Rhipicephalus* ticks in the study are labelled in bold, by trees representing the L segment (encoding the polymerase), M segment (encoding the glycoproteins Gn and Gc) and S segment (encoding the nucleoprotein N). Samples representing Clade II (the Africa 2 lineage) are highlighted in peach. Samples highlighted in blue represent a divergent clade, most closely related to Clade V (the Europe 1 geographic lineage). Sample tips are labelled by country of origin and accession number. The source of samples obtained from Genbank files (human, tick, sheep or unknown) are labelled as shown in the key.

We also describe for the first time, Macira virus; a novel member of the *Nairoviridae* detected in one pool of *R. appendiculatus* ticks collected from an environmental sample. This segmented virus was most closely related to Nayun tick virus, based on the S segment, but with an amino acid uncorrected pairwise distance of 33% (**Fig 4a**). The partial L segment for the virus had a distance of 49% from other orthonairoviruses. Species demarcation of the *Nairoviridae* is based on the rules governing CCHFV sequences, which have a p-distance of less than 6.8% in the polymerase protein (11).

Within the *Orthomyxoviridae*, we detected the *Quaranjavirus* Zambezi tick virus 1 (previously identified in ticks in Mozambique (12)) in 19 *R. appendiculatus* pools, including P44 from Arua district represented in **Fig 4b** representing a subspecies of this virus circulating in Uganda. We obtained genome coverage across five segments (PB1, PB2, PA, HA, NP) with an uncorrected p-distance of 2.2% in the PB1 protein with Zambezi tick virus 1 and 10-25% in other proteins with other *Quaranjavirus* species. Thogoto virus was also detected in one *Amblyomma* tick pool.

We detected the segmented Jingmen tick virus in 4 pools of *Rhipicephalus* ticks. In addition, four pools (P30, P47, P51 and P56) were found to contain the genome of a novel virus member of the segmented *Flaviviridae* family, with homology to the *Jingmen tick virus*, that we have named Engoha virus, all derived from *Rhipicephalus* ticks collected from both cattle and goats (**Fig 4c**) (13). Four segments representing the NS5-like, glycoprotein, NS3-like, capsid and membrane proteins were obtained by mapping to the closest reference sequences. Member viruses in this clade differ from each other in the NS5-like protein by 0.3-4%. We detected three divergent strains of Engoha virus with an uncorrected p-distance for the NS5-like protein of 2-3% and 3.1% from the nearest Mogiana tick virus reference sequence. We have named these as Engoha virus isolates 1-3. We also detected Trinbago virus, another member of the *Flaviviridae* in six *Rhipicephalus* tick pools from Nakaseke and Lyantonde districts, four of which were used for phylogenetic analysis (P158, P171, P172, P176; **Fig 4d**), a virus first detected in Trinidad and Tobago in *R. sanguineus*, *R. microplus*, and *A. ovale* ticks in 2019 and more recently detected in Kenya (14). Viruses in this clade are assigned to the *Pestivirus*- like group and species differ by 15% in the polyprotein amino acid sequence. The Ugandan sequence had an uncorrected p-distance of 5.4% and was confirmed as a Ugandan strain of Trinbago virus (13). Also within the *Flaviviridae*, we detected Bole tick virus 4 in 3 *Rhipicephalus* tick pools.

Within *R. appendiculatus* mature ticks collected from cattle in Nakaseke district (tick pool P214), we detected Barur virus within the *Ledantevirus* genus in the family *Rhabdoviridae* (Fig S3). This differed by 4.6% in the polymerase protein and 8.1% in the glycoprotein.

We detected Karukera tick virus in 16 *Amblyomma* (n=13) and *Rhipicephalus* (n=3) tick pools (15 from cattle and 1 from goats). This virus belongs to the *Mivirus* genus in the *Chuviridae* family previously reported from Guadeloupe in the Caribbean. We detected three segments representing the polymerase, glycoprotein and nucleoprotein. The polymerase from the a representative genome had an uncorrected p-distance of 3.2% compared with the Karukera reference genome, within the species demarcation criterion of 10% (15).

Finally, within the *Phenuiviridae* (**Fig 4f**), we detected three novel viruses in *Rhipicephalus* tick pools; Torwonik virus was detected in tick pool P29 (*Rhipicephalus* nymphs from Arua) and Enkwa virus in P194 (*Rhipicephalus* nymphs from Nakaseke) were placed within a *Bandavirus* clade that includes Forecariah virus, Bhanja virus, Razdan virus and Palma virus. These species differ in the polymerase by an uncorrected p-distance of 5% and 5.9% respectively from Bhanja virus. A more divergent species, named Ekuda virus, was detected in pool P196 (*R. appendiculatus* adult ticks from Nakaseke). We detected a fragment covering 18% of the polymerase which had an uncorrected p-distance of 37.3% to Kharabali tick virus polymerase, a *Phlebovirus*. Species demarcation for this family is proposed at 5% (16).

## Discussion

In this study, we aimed to investigate the diversity of ticks and of associated viruses in three distinct geographical areas of Uganda using semi-agnostic target enrichment NGS, a highly sensitive method that enables the detection of known and previously uncharacterized viruses. We detected six predominant species of ticks, collected from domestic animals and environmental collections belonging to the *Rhipicephalus, Amblyomma* and *Hyalomma* genera. *R. appendiculatus* ticks, the most prevalent species, are known to be widespread in Uganda and lack seasonal variation in abundance (17-20). They are also well-described vectors for zoonotic diseases, as they feed on a wide range of animals as well as humans (21, 22), including CCHFV (23, 24), Nairobi Sheep Disease Virus (NSDV) (25) and Kadam virus (26, 27), all of which are prevalent in Uganda.

Within the tick pools, we detected viruses belonging to 9 families (*Nairoviridae*, *Retroviridae*, *Orthomyxoviridae*, *Chuviridae*, *Rhabdoviridae*, *Phenuiviridae, Parvoviridae, Poxviridae* and *Flaviviridae*), 15 genera (*Alpharicinrhavirus, Bandavirus, Betaretrovirus, Ephemerovirus, Erythroparvovirus, Flavivirus, Jingmenvirus, Ledantevirus, Mivirus, Orthonairovirus, Parapoxvirus, Pestivirus, Phlebovirus, Quaranjavirus, Thogotovirus)* and 22 species, including 5 novel species (**Table 2**).

Notably, human and animal pathogens were frequently present within the *Nairoviridae*, including CCHFV (n=6), DUGV (n=16), NSDV (n=1) and unclassified (n=29), present in 53 tick pools. We detected a reassortant CCHFV variant containing segments L and S derived from Clade II (Africa 2 lineage) and a highly divergent M segment, most closely related to Clade V (Europe 1 lineage). These were detected within *R. appendiculatus* ticks in both Arua (TickP67) and Lyantonde (TickP143) districts. Variation in the M segment could be associated with immune escape as it encodes the glycoproteins that are the target of antibody-mediated responses to CCHFV (28). The presence of two highly divergent M lineages in Uganda could also be associated with variations in clinical presentation, immune response and species tropism and requires further investigation. Importantly, CCHFV was not restricted only to the known high-risk cattle corridor areas of Uganda but was also detected in Arua district, previously considered as a low-risk area. This finding indicates that CCHFV may be underestimated as a cause of acute illness in areas of Uganda outside the cattle corridor, a finding supported by a high seroprevalence in farmers and domestic animals in the same areas (29). We also detected 16 tick pools (derived from *Amblyomma* (n=5) and *Rhipicephalus* (n=11)), mostly concentrated in Arua district (n=15/16), that tested positive for Dugbe virus. Dugbe virus is infectious but does not normally cause disease in sheep and cattle and may cause a mild febrile illness in humans (30). Finally, a novel *Orthonairovirus* of unknown pathogenicity, that we named Macira virus was also detected in *R. appendiculatus* ticks from Arua. district. It was most closely related to Nayun tick virus recovered from *Rhipicephalus* ticks collected from Menglian district of Yunnan in China and Romania (22, 31) and is of unknown pathogenicity. Cross-reactivity and the potential for cross-competition among the nairoviruses described in this study is, as yet, undocumented.

The family *Orthomyxoviridae* was represented in 44 tick pools. Twenty-four contigs present in this family could not be reliably speciated due to short lengths, but were highly divergent, suggestive of novel species. The most frequently detected virus was Zambezi tick virus 1, present in 19 *Rhipicephalus* pools (16 tick pools derived from cattle, and 3 from goats), a virus in the genus *Quaranjavirus*, first identified in *Rhipicephalus* ticks in Mozambique in 2018 (12). We additionally detected Thogoto virus in *Amblyomma* ticks found on cattle. This species has been associated with infection in humans and may cause fatal disease, characterized by fever and neurological symptoms, including meningitis, alongside other members of the genus *Thogotovirus* (32, 33).

The family *Flaviviridae* were the third most common virus group represented in tick samples, and were detected in 21 tick pools. Trinbago virus was detected in 6 *Rhipicephalus* tick pools from Nakaseke, Lyantonde and Arua districts. This virus was reported in 2019 in Trinidad and Tobago in *Rhipicephalus* and less often *Amblyomma* ticks (13). We detected Jingmen tick virus (JMTV) in 4 tick pools. JMTV was first described in *R. microplus* ticks in China.

The virus has been described in Europe (it is also known as Alongshan virus in Russia and Finland) (20, 21), and more recently in Kenya in several tick species including *R. appendiculatus, R. evertsi evertsi, H. truncatum* collected from cattle and sheep, and *A.sparsum, A.nuttalli* and *Amblyoma* nymphs collected from tortoises (22). In Uganda, JMTV has also been isolated from non-human primates (colobus monkeys), highlighting the potential risk to primates (23). A probable human infection was described in Jingmen, China (17, 18), and it was also described as a coinfection with CCHFV in a fatal human case in Kosovo (19). We also describe a novel segmented *Flaviviridae*-like virus, that we have named Engoha virus, in four independent pools of *R. evertsi evertsi* (TickP30, TickP47, TickP51, TickP56) collected from cattle and goats in Arua and Lyantonde districts, sharing a clade with *Heilongjiang tick virus* (detected in *Dermacentor* ticks in China), Amblyomma virus, Jingmen tick virus, Mogiana tick virus, and Alongshan virus. We describe 3 strains of this virus, based on nucleotide diversity, most evident in the glycoprotein.

We detected 18 pools containing contigs of species in the *Chuviridae* (17 tick pools derived from cattle and 1 from goats). Two of these remained unassigned, as they were too short to classify as novel species, based on current speciation criteria. The majority of the pools (n=16) were positive for Karukera tick virus, 13 in *Amblyomma* tick pools and 3 in *Rhipicephalus* pools. Karukera tick virus was described recently in Guadeloupe and Martinique with a similar distribution in *Amblyomma* and *Rhipicephalus* ticks (34).

Twelve tick pools had detectable genomes derived from members of the *Rhabdoviridae*. Six of these were highly divergent, but were short contigs and speciation criteria could not be applied. Wuhan tick virus 1 was present in three pools, *Rhipicephalus* associated rhabdo-like virus in two and Barur virus, a member of the *Ledantevirus* genus in one, all of which have been described previously in ticks and which have not been associated to date with human infection.

We detected 5 tick pools infected with virus species belonging to the family *Phenuiviridae*, three of which were novel viruses (newly described as Enkuda virus, Enkwa virus and Torwonik virus; Figure 3). The family *Phenuiviridae* contains several arboviruses that can replicate in insect and mammalian hosts, including Rift Valley fever virus (RVFV) and Severe Fever with Thrombocytopenia syndrome virus (SFTSV) (35). Torwonik and Enkwa viruses are situated phylogenetically within the *Bhanja* serogroup clade containing Bhanja virus (BHAV), Forecariah virus (FORV), Kismayo virus (KISV), and Palma virus (PALV) (36). BHAV has previously been isolated from *R. appendiculatus* ticks in Kenya and Nigeria (26, 29). The medical implication of BHAV infection has not been documented in East Africa, but in the European region and western part of Africa, BHAV has been linked to human illness (30). These viruses therefore require further investigation regarding their ability to infect mammals and to cause disease.

We detected several viruses not known to be vectored by ticks that may derive from the blood meal of the tick host, including Enzootic nasal tumor virus of goats belonging to the genus *Betaretrovirus* in the family *Retroviridae* in 5 tick pools, all from goats, a retrovirus associated with infectious nasal adenocarcinoma in this species (ENTV-2). We also detected Bovine parvovirus 3 (BPV) in a single sample from *Amblyomma* ticks derived from cattle.

BPV may cause respiratory and gastrointestinal disease in cows and belongs to the *Bocaparvovirus* genus of the *Parvoviridae* family. Finally, we detected *Pseudocowpoxvirus*, (family *Poxviridae* and genus *Parapoxvirus*), also in a single pool of *Amblyomma* ticks derived from cattle. This virus can infect both livestock causing bovine popular stomatitis and humans, in whom it may cause a vesicular rash and fever.

## Conclusions

In this study, we describe the diversity of viruses detected in ticks in Uganda. We detected 9 families, 15 genera and 22 species of virus, including 8 species known to cause disease in animals and/or humans and 5 novel species of unknown pathogenic potential. Further studies to investigate the role of the novel viruses detected in this study in causing animal or human disease is indicated.

## Materials and Methods

### Ethics

This study was undertaken as part of the ArboViral Infection study (AVI) approved by the Research Ethics Review Committee at the College of Veterinary Medicine, Animal Resources and Biosecurity of Makerere University, Kampala, Uganda (Reference Number: SVARREC/20/2018) and by the Uganda National Council for Science and Technology (Reference Number: HS 2485). Informed written consent was obtained from farmers, before ticks were collected from livestock.

### Study design

We conducted a cross-sectional study in three districts of Uganda between December 2018 and August 2019. We selected two areas within the cattle corridor (Nakaseke and Lyantonde), classified as high risk districts because of their location in an area of high cattle density and previously-reported human cases of CCHF in the area (Fig 1). We also selected an area (Arua district) outside the cattle corridor (24, 29, 37, 38), classified as a low-risk area where CCHF had never previously been reported, as a control site. The study in Lyantonde coincided with a CCHF outbreak, and ecological sampling of ticks for virome analysis was undertaken as part of an outbreak investigation (39).

### Selection of study area and the farms

A multistage probability sampling approach was used in which one sub-county was randomly selected from the available list of sub-counties in the districts of Arua and Nakaseke.

Thereafter, a randomized selection of households that kept livestock within the selected sub-counties was made to reach a predetermined total of 1,215 livestock samples (40). In Lyantonde, farms were selected based on a trace back of a fatal case of CCHF in the district, to identify CCHFV spillover farm(s) associated with the outbreak (39). Tick samples were collected from a minimum of 25% of randomly selected animals on each farm.

### Tick collection and processing

Livestock ticks were collected from half of the body of every selected animal as previously described (29). Ticks were transported in 70% ethanol for identification to species level using morphological keys (19, 41) under a stereo-microscope (Stereo Discovery V12, Zeiss, Birkerød, Denmark). Tick pools containing 5-10 adult ticks, 10-20 nymphs and 25 larvae were made according to their collection sites, species, sex, and the host animal. All tick pools were then crushed in 0·5ml of Agencourt RNAdvance lysis buffer in a Genogrinder 2000 (OPS Diagnostics, Lebanon, NJ, USA), followed by downstream RNA extraction procedures as per manufacturer’s instructions including a DNase treatment step at 37°C for 15 min (Beckman Coulter).

### Detection of viruses in tick samples

Tick pools were investigated for the presence of viral genomes using target enrichment NGS as previously described, using customised ArboCap enrichment probes targeting all arboviruses (Agilent) (42). Viral genomes were detected following *de novo* assembly using SPADES and IDBA-UD followed by BLASTn and mapping to the relevant viral reference sequences. Maximum likelihood phylogenetic analysis was carried out using IQTREE 1.6.12 (43) with the best substitution model (ModelFinder) (44) and 1000 ultra-fast bootstrap replicates (45). Uncorrected nucleotide and amino acid pairwise distances were calculated on MAFFT alignments using MEGA-X. We followed guidance from the ICTV and relevant literature for the classification of novel viruses by family, genus and species. Recommended criteria used are summarised in **Supplementary Table 2**.

### Viral diversity estimates

Virus hits were aggregated by the genus of the closest reference sequence. We used the estimate-richness function of the R phyloseq package to compute alpha diversity (richness and Simpson’s diversity index) of virus genera across sampling districts and tick species. The distribution of alpha diversity was compared amongst sampling locations and tick species using the Kruskal-Wallis test. Multiple pairwise comparisons of alpha diversity amongst tick species were carried out using Dunn’s test with a Bonferroni correction. Differences in tick pool positivity of virus genera between sampling sites was compared using Fisher exact test with a Bonferroni correction. Analysis was carried out in R version 4.2.1.

## Author contributions

SAA, RT, CM, and ECT conceived the idea and designed the study, collected data, drafted the manuscript

LT, AF, SA, ARA, PV, WN, TN, JN, JJL and SAA supported in in field sample collection and analysis and sequencing analysis.

SA, MN, AS, MNH, JGS, ECT and SAA contributed in data analysis and reporting.

ECT, AF, JJL, PK, CM, RT, JE, SO, DM provided supervision of the project and writing.

All authors had full access to all the data of the study and had final responsibility for the decision to submit the final manuscript for publication

## Acknowledgements

We thank the study participants for making this study possible by providing samples and answering the questionnaire. We acknowledge support from the Department of Arbovirology, Emerging and Re-emerging infectious Diseases (UVRI), National Animal Diseases Diagnostics and Epidemiology Centre (MAAIF), MRC-University of Glasgow, Centre for Virus Research (UoG), and Makerere University School of Veterinary Medicine and Animal Resources.

---

**Supplementary Figure 1:**
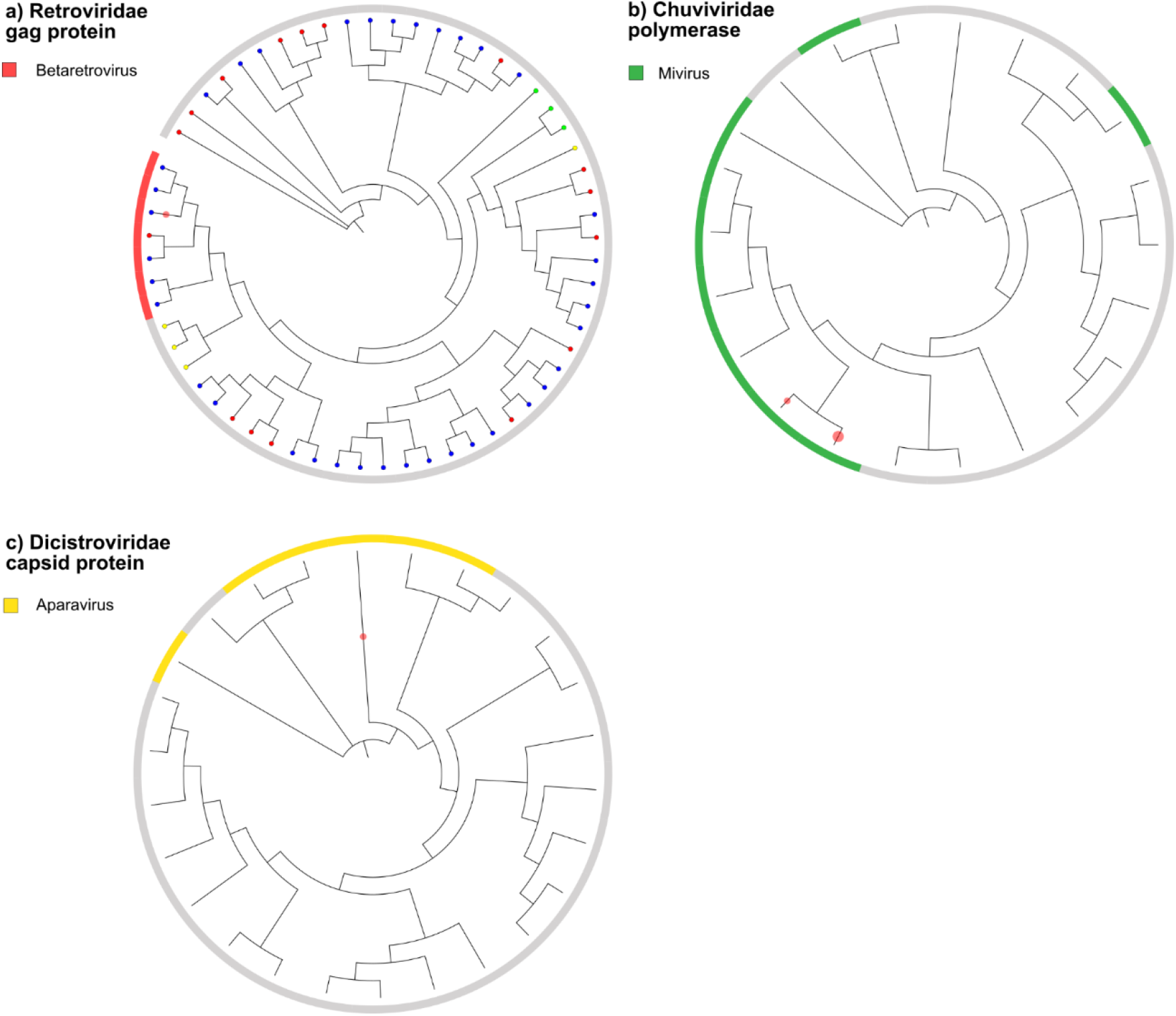
EPA-NG placement of additional contigs. Contigs were placed within the (a) *Retroviridae*, (b) *Chuviridae* and (c) *Dicistroviridae*

**Supplementary Figure 2:**
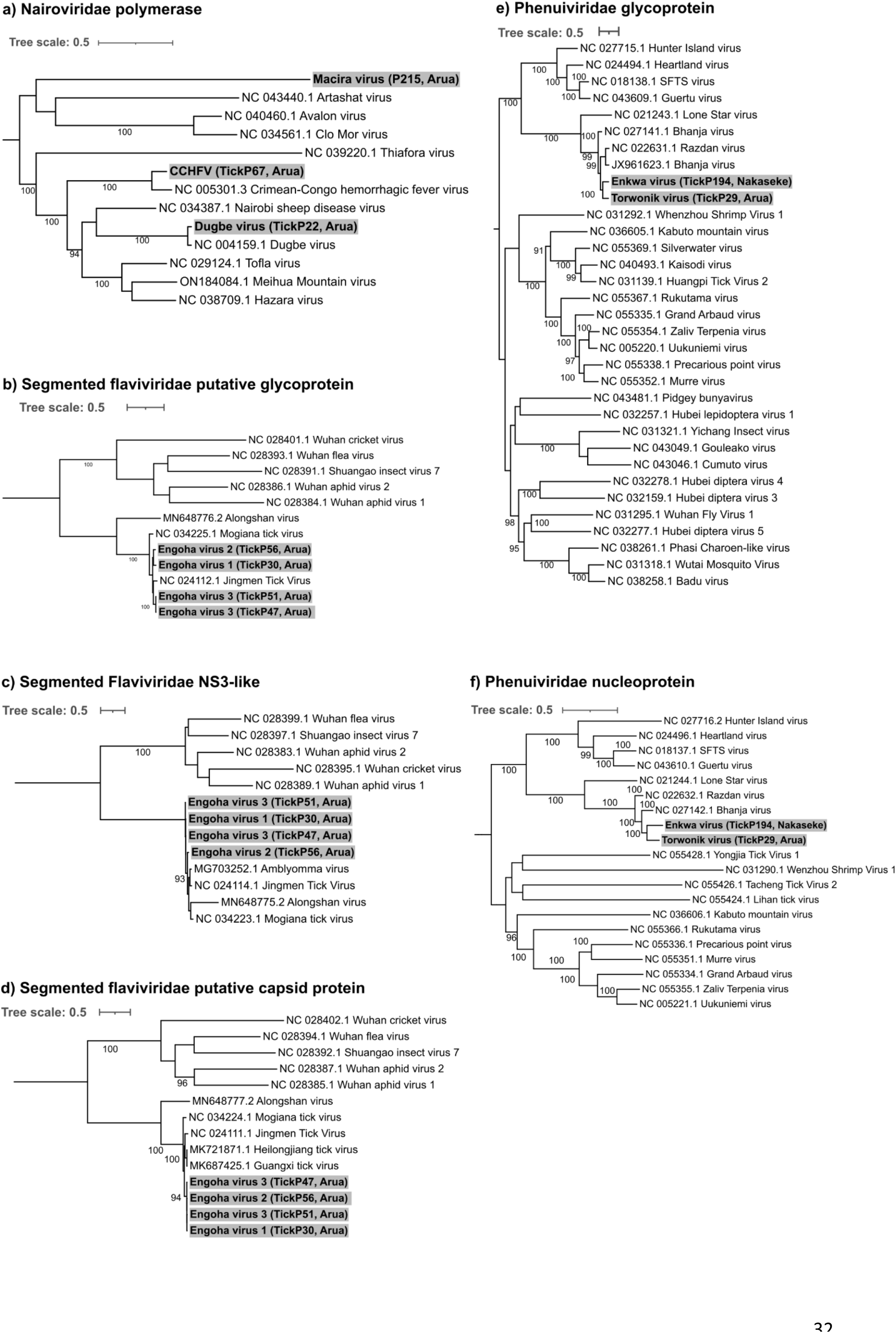
Maximum likelihood phylogenetic analysis of additional contigs (a) *Nairoviridae* polymerase (b) Segmented *Flaviviridae* glycoprotein (c) Segmented *Flaviridae* NS3-like protein (d) Segmented *Flaviridae* capsid protein (e) *Phenuiviridae* glycoprotein and (f) *Phenuiviridae* nucleoprotein

**Supplementary Figure 3:**
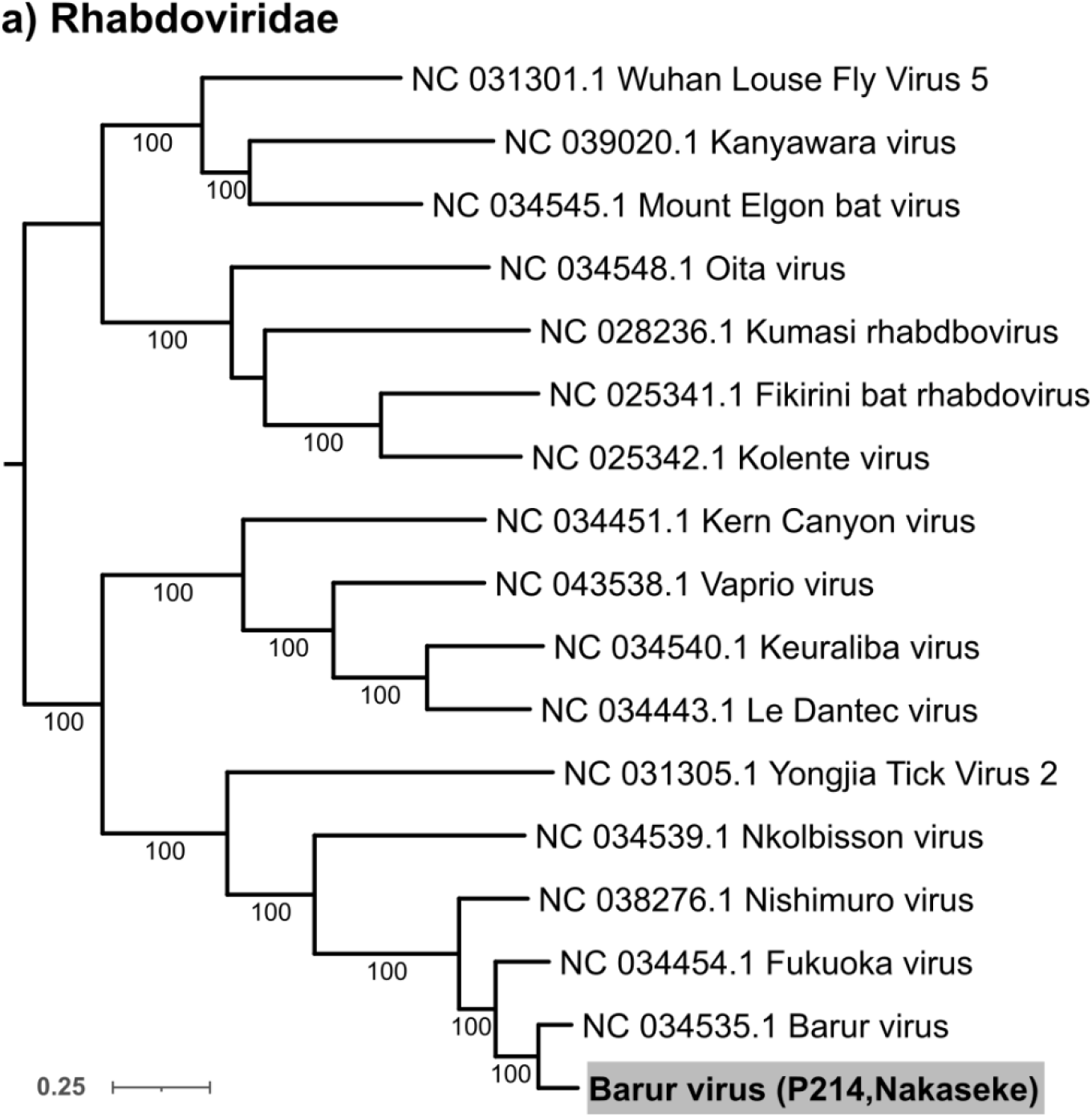
Representative maximum likelihood phylogenetic analysis of the *Rhabdoviridae*

## Supporting information captions

**Supplementary Table 1.**
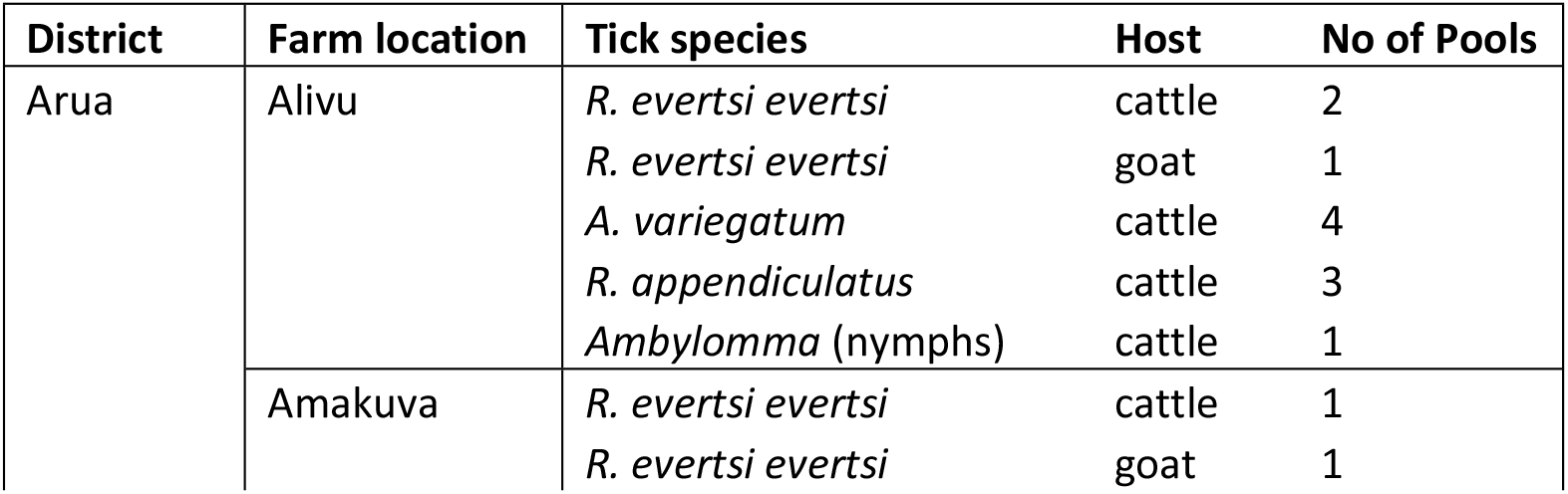
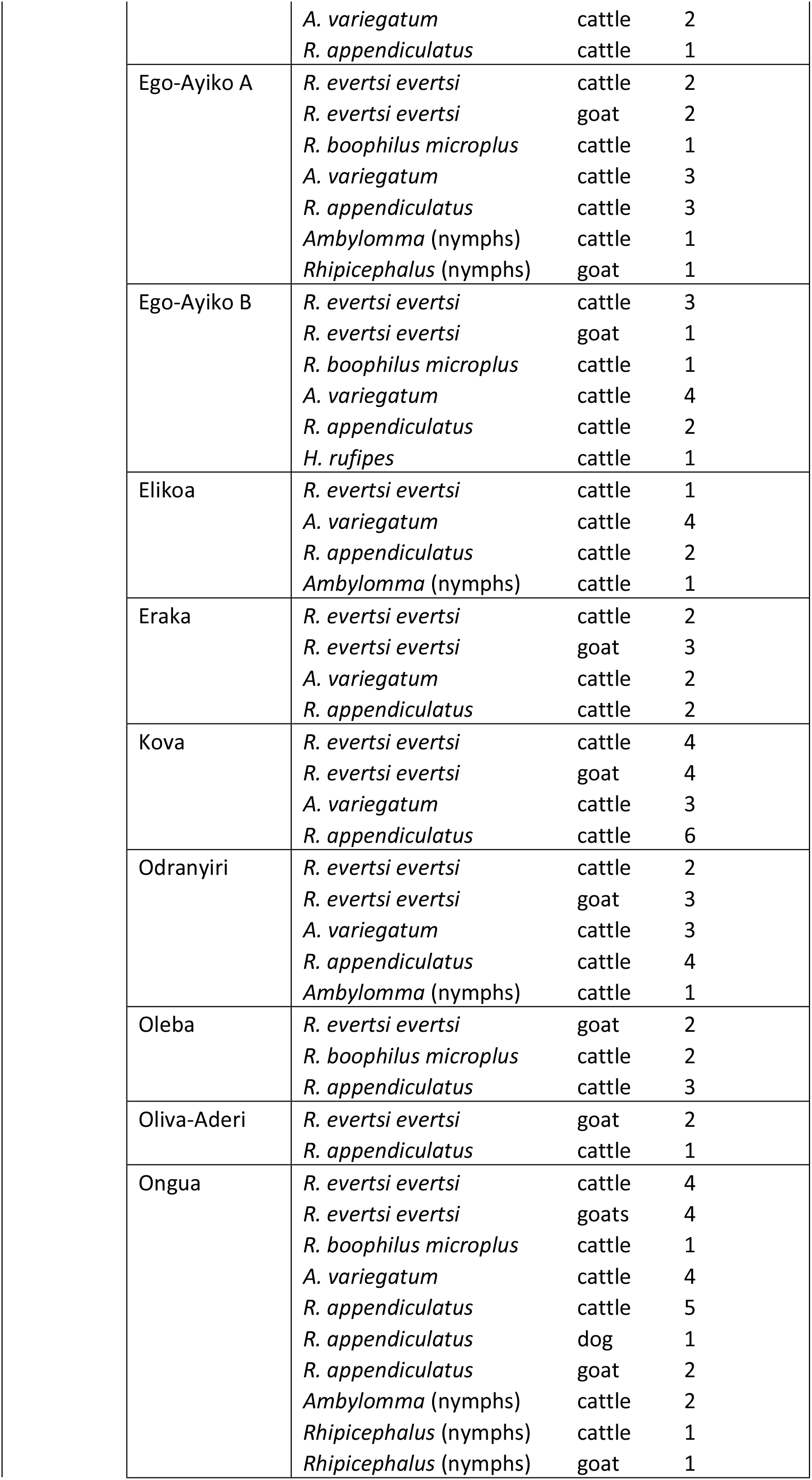
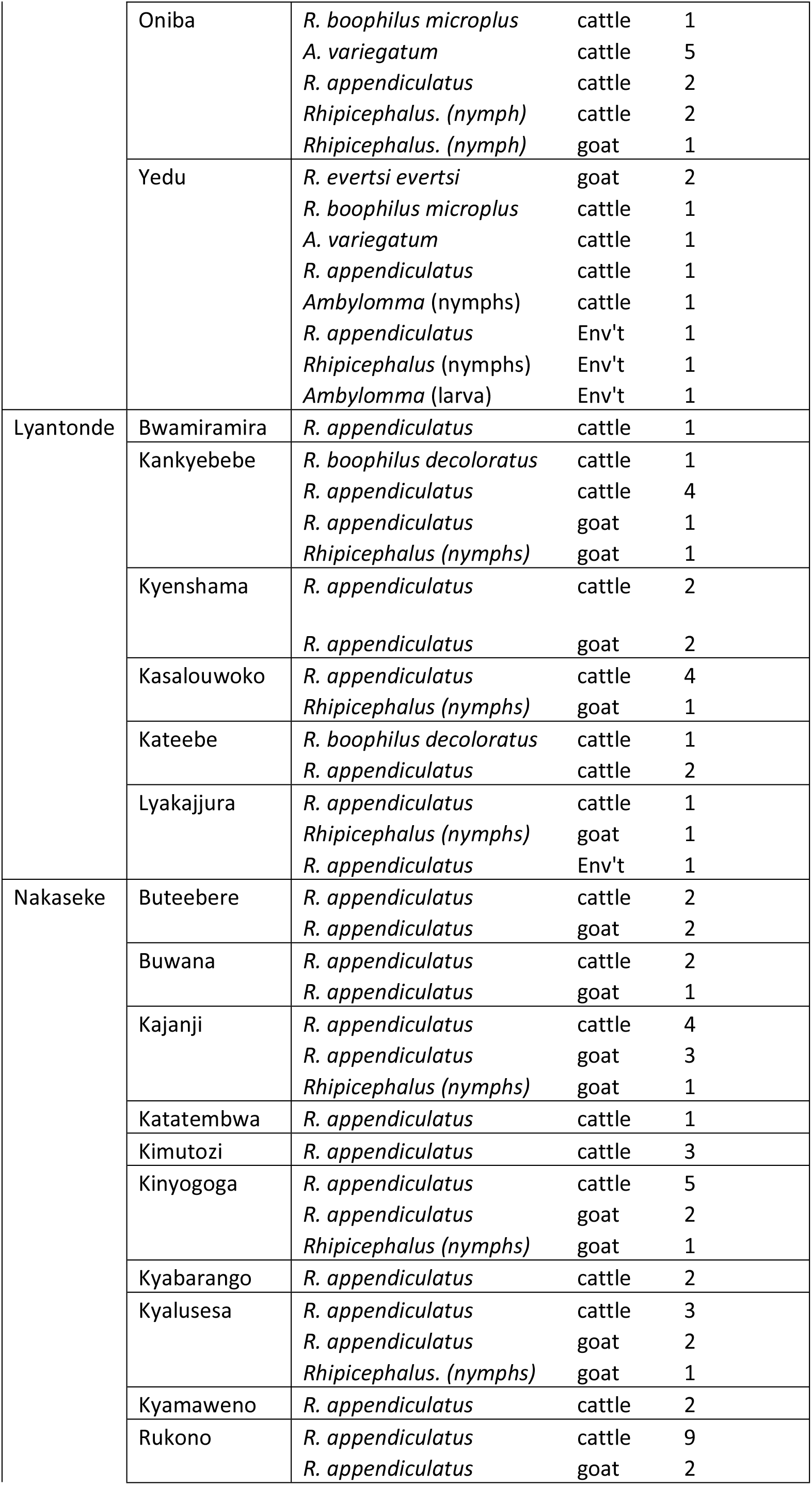
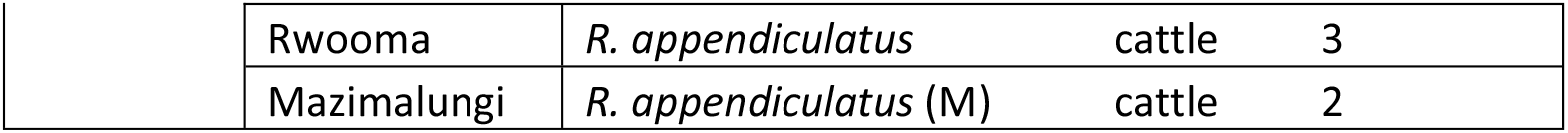
Location of tick species sampling by host and number of pools

| District | Farm location | Tick species | Host | No of Pools |
| --- | --- | --- | --- | --- |
| Arua | Alivu | *R. evertsi evertsi* | cattle | 2 |
|  |  | *R. evertsi evertsi* | goat | 1 |
|  |  | *A. variegatum* | cattle | 4 |
|  |  | *R. appendiculatus* | cattle | 3 |
|  |  | *Ambylomma* (nymphs) | cattle | 1 |
|  | Amakuva | *R. evertsi evertsi* | cattle | 1 |
|  |  | *R. evertsi evertsi* | goat | 1 |

|  |  |  |  |
| --- | --- | --- | --- |
|  | *A. variegatum* | cattle | 2 |
|  | *R. appendiculatus* | cattle | 1 |
| Ego-Ayiko A | *R. evertsi evertsi* | cattle | 2 |
|  | *R. evertsi evertsi* | goat | 2 |
|  | *R. boophilus microplus* | cattle | 1 |
|  | *A. variegatum* | cattle | 3 |
|  | *R. appendiculatus* | cattle | 3 |
|  | *Ambylomma* (nymphs) | cattle | 1 |
|  | *Rhipicephalus* (nymphs) | goat | 1 |
| Ego-Ayiko B | *R. evertsi evertsi* | cattle | 3 |
|  | *R. evertsi evertsi* | goat | 1 |
|  | *R. boophilus microplus* | cattle | 1 |
|  | *A. variegatum* | cattle | 4 |
|  | *R. appendiculatus* | cattle | 2 |
|  | *H. rufipes* | cattle | 1 |
| Elikoa | *R. evertsi evertsi* | cattle | 1 |
|  | *A. variegatum* | cattle | 4 |
|  | *R. appendiculatus* | cattle | 2 |
|  | *Ambylomma* (nymphs) | cattle | 1 |
| Eraka | *R. evertsi evertsi* | cattle | 2 |
|  | *R. evertsi evertsi* | goat | 3 |
|  | *A. variegatum* | cattle | 2 |
|  | *R. appendiculatus* | cattle | 2 |
| Kova | *R. evertsi evertsi* | cattle | 4 |
|  | *R. evertsi evertsi* | goat | 4 |
|  | *A. variegatum* | cattle | 3 |
|  | *R. appendiculatus* | cattle | 6 |
| Odranyiri | *R. evertsi evertsi* | cattle | 2 |
|  | *R. evertsi evertsi* | goat | 3 |
|  | *A. variegatum* | cattle | 3 |
|  | *R. appendiculatus* | cattle | 4 |
|  | *Ambylomma* (nymphs) | cattle | 1 |
| Oleba | *R. evertsi evertsi* | goat | 2 |
|  | *R. boophilus microplus* | cattle | 2 |
|  | *R. appendiculatus* | cattle | 3 |
| Oliva-Aderi | *R. evertsi evertsi* | goat | 2 |
|  | *R. appendiculatus* | cattle | 1 |
| Ongua | *R. evertsi evertsi* | cattle | 4 |
|  | *R. evertsi evertsi* | goats | 4 |
|  | *R. boophilus microplus* | cattle | 1 |
|  | *A. variegatum* | cattle | 4 |
|  | *R. appendiculatus* | cattle | 5 |
|  | *R. appendiculatus* | dog | 1 |
|  | *R. appendiculatus* | goat | 2 |
|  | *Ambylomma* (nymphs) | cattle | 2 |
|  | *Rhipicephalus* (nymphs) | cattle | 1 |
|  | *Rhipicephalus* (nymphs) | goat | 1 |
|  | Oniba | *R. boophilus microplus* | cattle | 1 |
|  |  | *A. variegatum* | cattle | 5 |
|  |  | *R. appendiculatus* | cattle | 2 |
|  |  | *Rhipicephalus. (nymph)* | cattle | 2 |
|  |  | *Rhipicephalus. (nymph)* | goat | 1 |
|  | Yedu | *R. evertsi evertsi* | goat | 2 |
|  |  | *R. boophilus microplus* | cattle | 1 |
|  |  | *A. variegatum* | cattle | 1 |
|  |  | *R. appendiculatus* | cattle | 1 |
|  |  | *Ambylomma (nymphs)* | cattle | 1 |
|  |  | *R. appendiculatus* | Env't | 1 |
|  |  | *Rhipicephalus (nymphs)* | Env't | 1 |
|  |  | *Ambylomma (larva)* | Env't | 1 |
| Lyantonde | Bwamiramira | *R. appendiculatus* | cattle | 1 |
|  | Kankyebebe | *R. boophilus decoloratus* | cattle | 1 |
|  |  | *R. appendiculatus* | cattle | 4 |
|  |  | *R. appendiculatus* | goat | 1 |
|  |  | *Rhipicephalus (nymphs)* | goat | 1 |
|  | Kyenshama | *R. appendiculatus* | cattle | 2 |
|  |  | *R. appendiculatus* | goat | 2 |
|  | Kasalouwoko | *R. appendiculatus* | cattle | 4 |
|  |  | *Rhipicephalus (nymphs)* | goat | 1 |
|  | Kateebe | *R. boophilus decoloratus* | cattle | 1 |
|  |  | *R. appendiculatus* | cattle | 2 |
|  | Lyakajjura | *R. appendiculatus* | cattle | 1 |
|  |  | *Rhipicephalus (nymphs)* | goat | 1 |
|  |  | *R. appendiculatus* | Env't | 1 |
| Nakaseke | Buteebere | *R. appendiculatus* | cattle | 2 |
|  |  | *R. appendiculatus* | goat | 2 |
|  | Buwana | *R. appendiculatus* | cattle | 2 |
|  |  | *R. appendiculatus* | goat | 1 |
|  | Kajanji | *R. appendiculatus* | cattle | 4 |
|  |  | *R. appendiculatus* | goat | 3 |
|  |  | *Rhipicephalus (nymphs)* | goat | 1 |
|  | Katatembwa | *R. appendiculatus* | cattle | 1 |
|  | Kimutozi | *R. appendiculatus* | cattle | 3 |
|  | Kinyogoga | *R. appendiculatus* | cattle | 5 |
|  |  | *R. appendiculatus* | goat | 2 |
|  |  | *Rhipicephalus (nymphs)* | goat | 1 |
|  | Kyabarango | *R. appendiculatus* | cattle | 2 |
|  | Kyalusesa | *R. appendiculatus* | cattle | 3 |
|  |  | *R. appendiculatus* | goat | 2 |
|  |  | *Rhipicephalus. (nymphs)* | goat | 1 |
|  | Kyamaweno | *R. appendiculatus* | cattle | 2 |
|  | Rukono | *R. appendiculatus* | cattle | 9 |
|  |  | *R. appendiculatus* | goat | 2 |

|  |  |  |  |
| --- | --- | --- | --- |
| Rwooma | *R. appendiculatus* | cattle | 3 |
| Mazimalungi | *R. appendiculatus* (M) | cattle | 2 |

**Supplementary Table 2.**
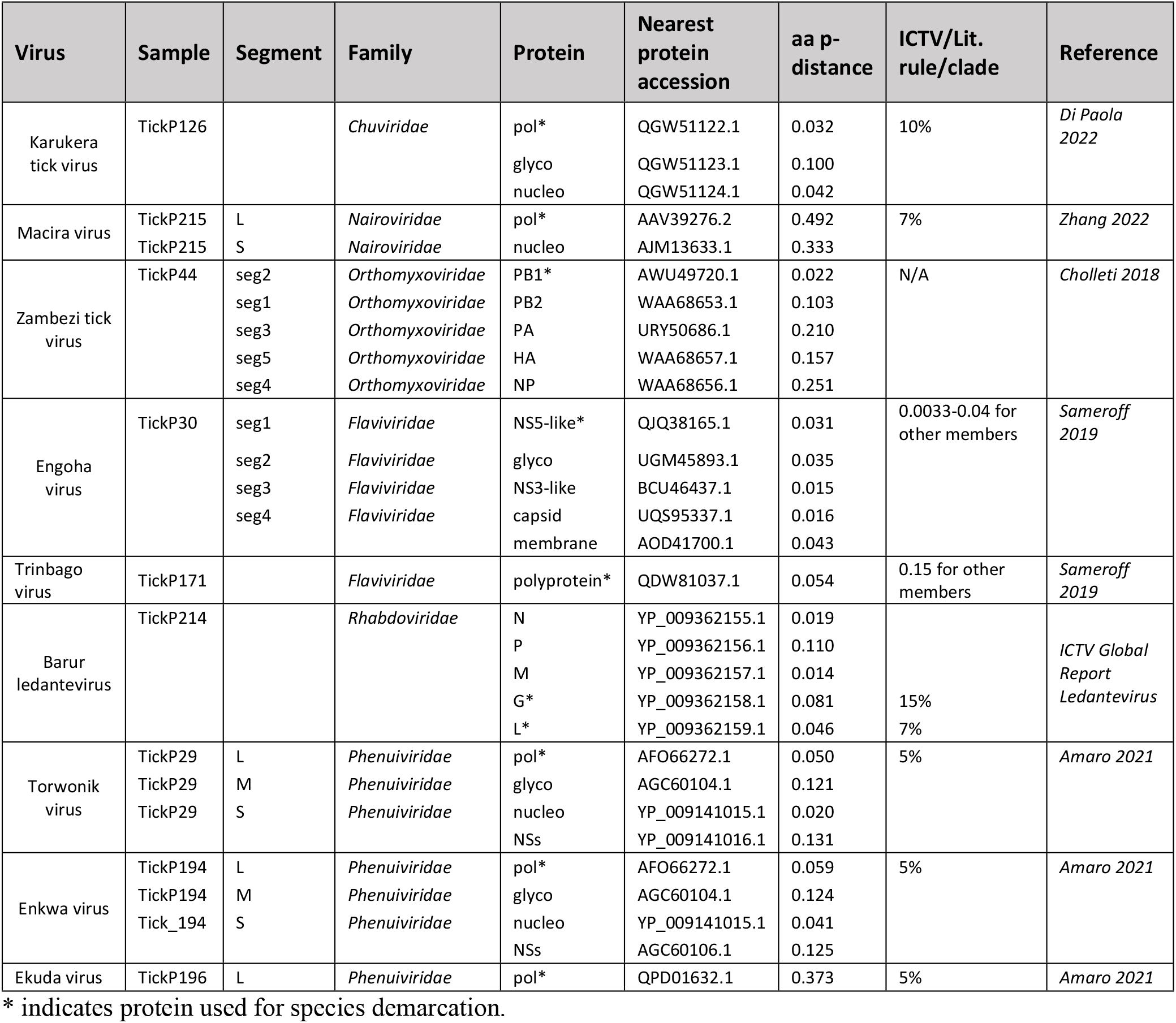
Criteria used to classify novel viruses by family

**Supplementary Table 3:**
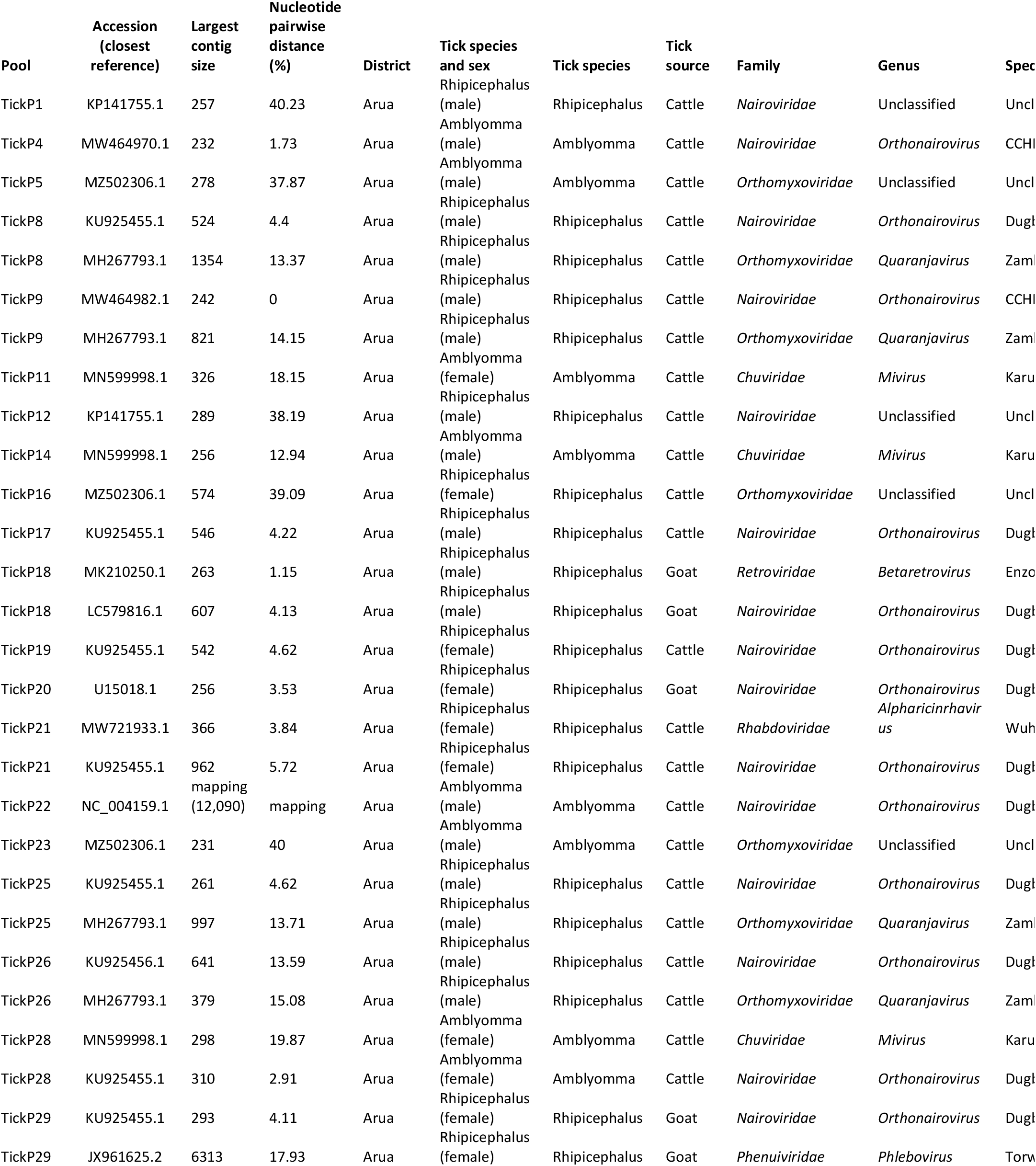
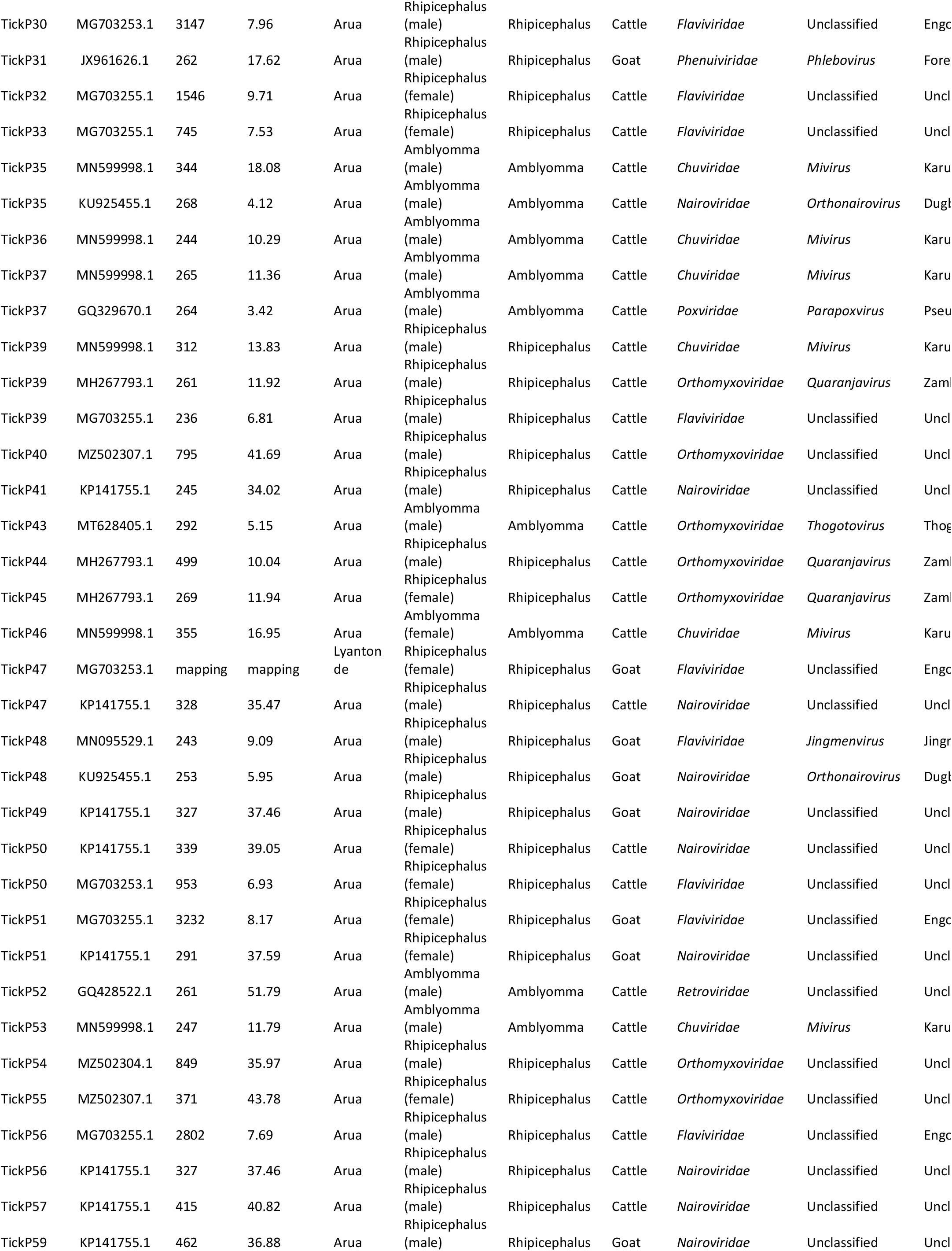
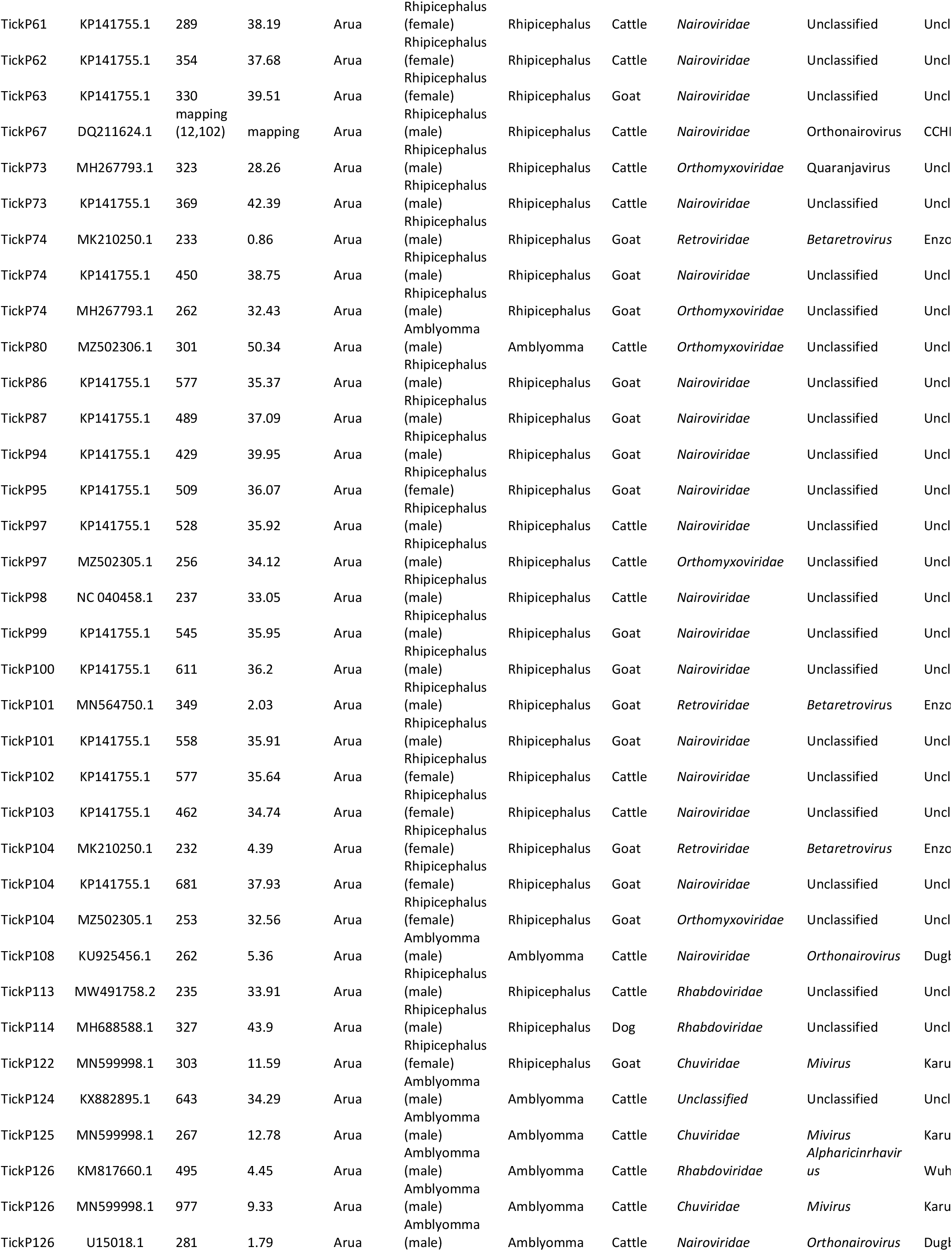
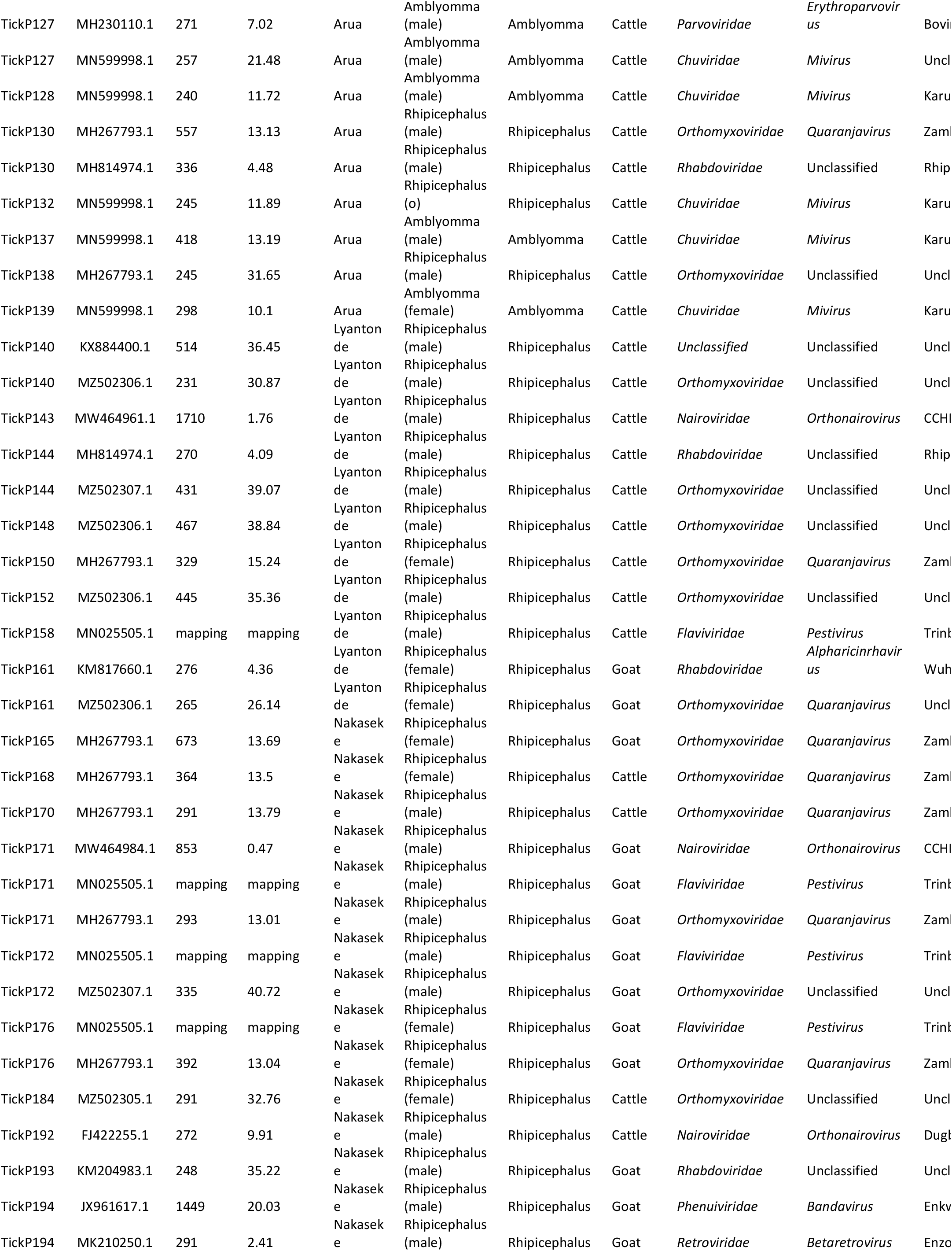
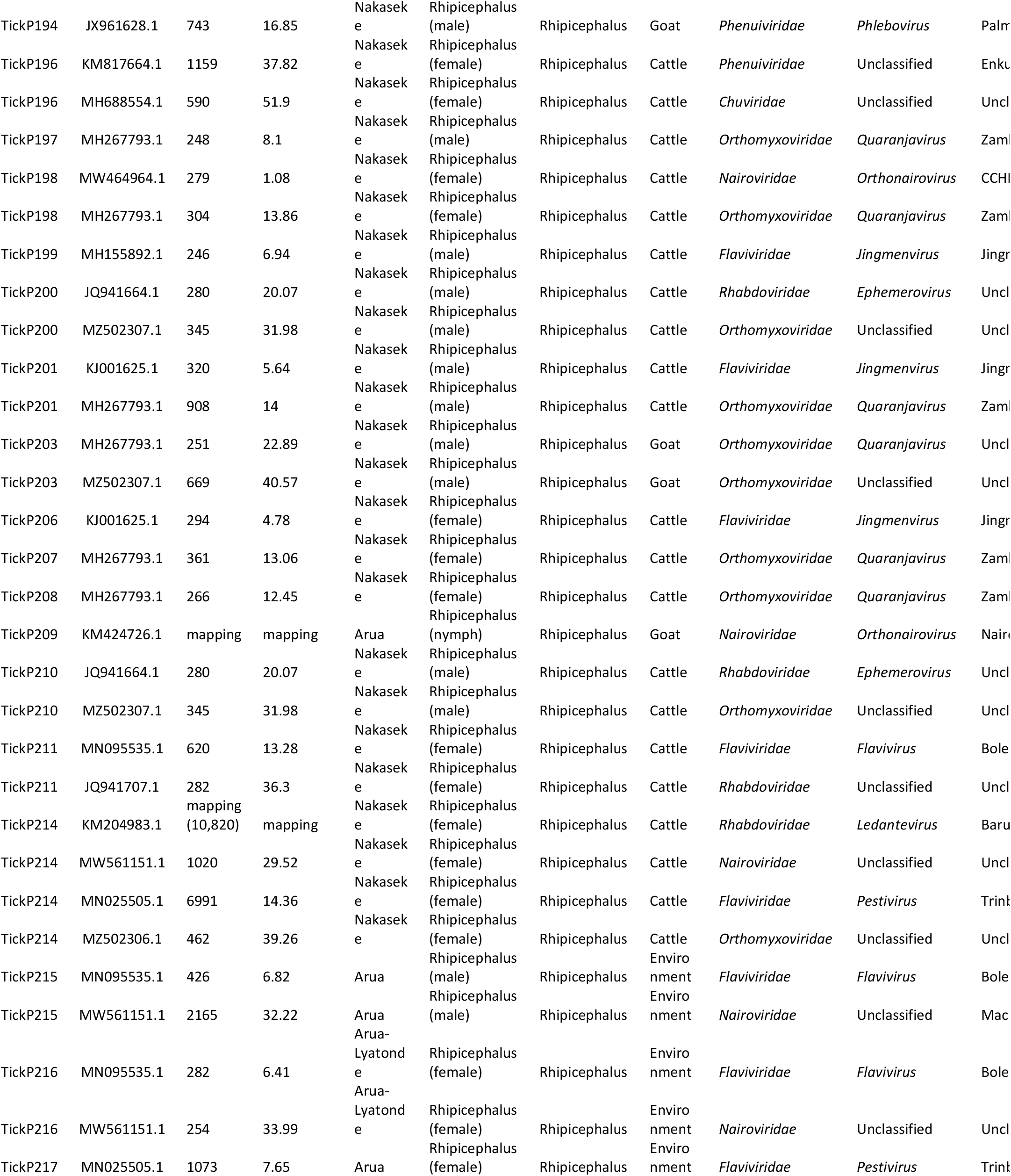
Tick pools by virus detected, contig size and nucleotide pairwise distance

**Supplementary Table 4.** Positivity rate per tick pool (by genus) between sampling sites

| Genus | Group 1 | Group 2 | p-value* | p.adjusted | P.adj.symbol |
| --- | --- | --- | --- | --- | --- |
| *Orthonairovirus* | Arua | Lyantonde | 0.0398 | 0.12 | ns |
| *Orthonairovirus* | Arua | Nakaseke | 0.00113 | 0.003 | ** |
| *Orthonairovirus* | Lyantonde | Nakaseke | 1 | 10.948 | ns |
| *Quaranjavirus* | Arua | Lyantonde | 0.00764 | 0.02 | * |
| *Quaranjavirus* | Arua | Nakaseke | 0.000338 | 0.001 | ** |
| *Quaranjavirus* | Lyantonde | Nakaseke | 1 | 10.948 | ns |

\*Samples were tested by pairwise Fisher test with p-values adjusted using Bonferroni method

